# Relevance of SARS-CoV-2 related factors ACE2 and TMPRSS2 expressions in gastrointestinal tissue with pathogenesis of digestive symptoms, diabetes-associated mortality, and disease recurrence in COVID-19 patients

**DOI:** 10.1101/2020.04.14.040204

**Authors:** Ashutosh Kumar, Muneeb A. Faiq, Vikas Pareek, Khursheed Raza, Ravi K. Narayan, Pranav Prasoon, Pavan Kumar, Maheswari Kulandhasamy, Chiman Kumari, Kamla Kant, Himanshu N. Singh, Rizwana Qadri, Sada N. Pandey, Santosh Kumar

**Affiliations:** Etiologically Elusive Disorders Research Network (EEDRN), New Delhi, India; Department of Anatomy, All India Institute of Medical Sciences (AIIMS), Patna, India; New York University (NYU) Langone Health Center, NYU Robert I Grossman School of Medicine, New York, New York, USA; National Brain Research Center, Manesar, Haryana, India; Department of Anatomy, All India Institute of Medical Sciences, Deoghar, India; Pittsburgh Center for Pain Research, School of Medicine, University of Pittsburgh, Pittsburgh, Pennsylvania, USA; Department of Pediatrics, Medical University of South Carolina, Charleston, USA; Department of Biochemistry, Maulana Azad Medical College (MAMC), New Delhi, India; Department of Anatomy, Postgraduate Institute of Medical Education and Research (PGIMER), Chandigarh, India; Department of Microbiology, All India Institute of Medical Sciences (AIIMS), Bathinda, India; TAGC-INSERM, U1090, Aix Marseille University, Marseille, France; Neuro-oncology Laboratory, Rockefeller University, New York, New York, USA; Department of Zoology, Banaras Hindu University (BHU), Varanasi, India; Department of Anesthesiology and Critical Care Medicine, School of Medicine, Johns Hopkins University, Baltimore, USA

**Keywords:** SARS-CoV2, digestive symptoms, recurrence, amino acid transporter, glucose transporter

## Abstract

**Introduction:** COVID-19 is caused by a new strain of coronavirus called SARS-coronavirus-2 (SARS-CoV-2), which is a positive sense single strand RNA virus. In humans, it binds to angiotensin converting enzyme 2 (ACE2) with the help a structural protein on its surface called the S-spike. Further, cleavage of the viral spike protein (S) by the proteases like transmembrane serine protease 2 (TMPRSS2) or Cathepsin L (CTSL) is essential to effectuate host cell membrane fusion and virus infectivity. COVID-19 poses intriguing issues with imperative relevance to clinicians. The pathogenesis of GI symptoms, diabetes-associated mortality, and disease recurrence in COVID-19 are of particular relevance because they cannot be sufficiently explained from the existing knowledge of the viral diseases. Tissue specific variations of SARS-CoV-2 cell entry related receptors expression in healthy individuals can help in understanding the pathophysiological basis the aforementioned collection of symptoms.

**Materials and Methods:** The data were downloaded from the Human Protein Atlas available at (https://www.proteinatlas.org/humanproteome/sars-cov-2) and the tissue specific expressions (both mRNA and protein) of ACE2 and TMPRSS2 as yielded from the studies with RNA sequencing and immunohistochemistry (IHC) were analyzed as a function of the various components of the digestive tract. A digestive system specific functional enrichment map of ACE2 gene was created using g:profiler (https://biit.cs.ut.ee/gprofiler/gost) utility and the data were visualized using Cytoscape software, version 3.7.2 (https://cytoscape.org/).

**Results:** The correlated expression (transcriptomic and proteomic) of ACE2 (to which SARS-CoV-2 binds through the S-spike) was found to be enriched in the lower gastrointestinal tract (GIT) (highest in small intestine, followed by colon and rectum), and was undetectable in the upper GIT components: mouth cavity (tongue, oral mucosa, and salivary glands), esophagus, and stomach. High expression of ACE2 was noted in the glandular cells as well as in the enterocytes in the lining epithelium (including brush border epithelium). Among other digestive system organs, Gall bladder (GB) showed high expression of ACE2 in glandular cells, while any protein expression was undetectable in liver and pancreas. TMPRSS2 was found enhanced in GIT and exocrine glands of pancreas, and co-localized with ACE2 in enterocytes.

**Conclusions:** Based on the findings of this study and supportive evidence from the literature we propose that a SARS-CoV-2 binding with ACE2 mediates dysregulation of the sodium dependent nutrient transporters and hence may be a plausible basis for the digestive symptoms in COVID-19 patients. ACE2 mediated dysregulation of sodium dependent glucose transporter (SGLT1 or SLC5A1) in the intestinal epithelium also links it to the pathogenesis of diabetes mellitus which can be a possible reason for the associated mortality in COVID-19 patients with diabetes. High expression of ACE2 in mucosal cells of the intestine and GB make these organs potential sites for the virus entry and replication. Continued replication of the virus at these ACE2 enriched sites may be a basis for the disease recurrence reported in some, thought to be cured, patients.

**Graphical Abstract:** 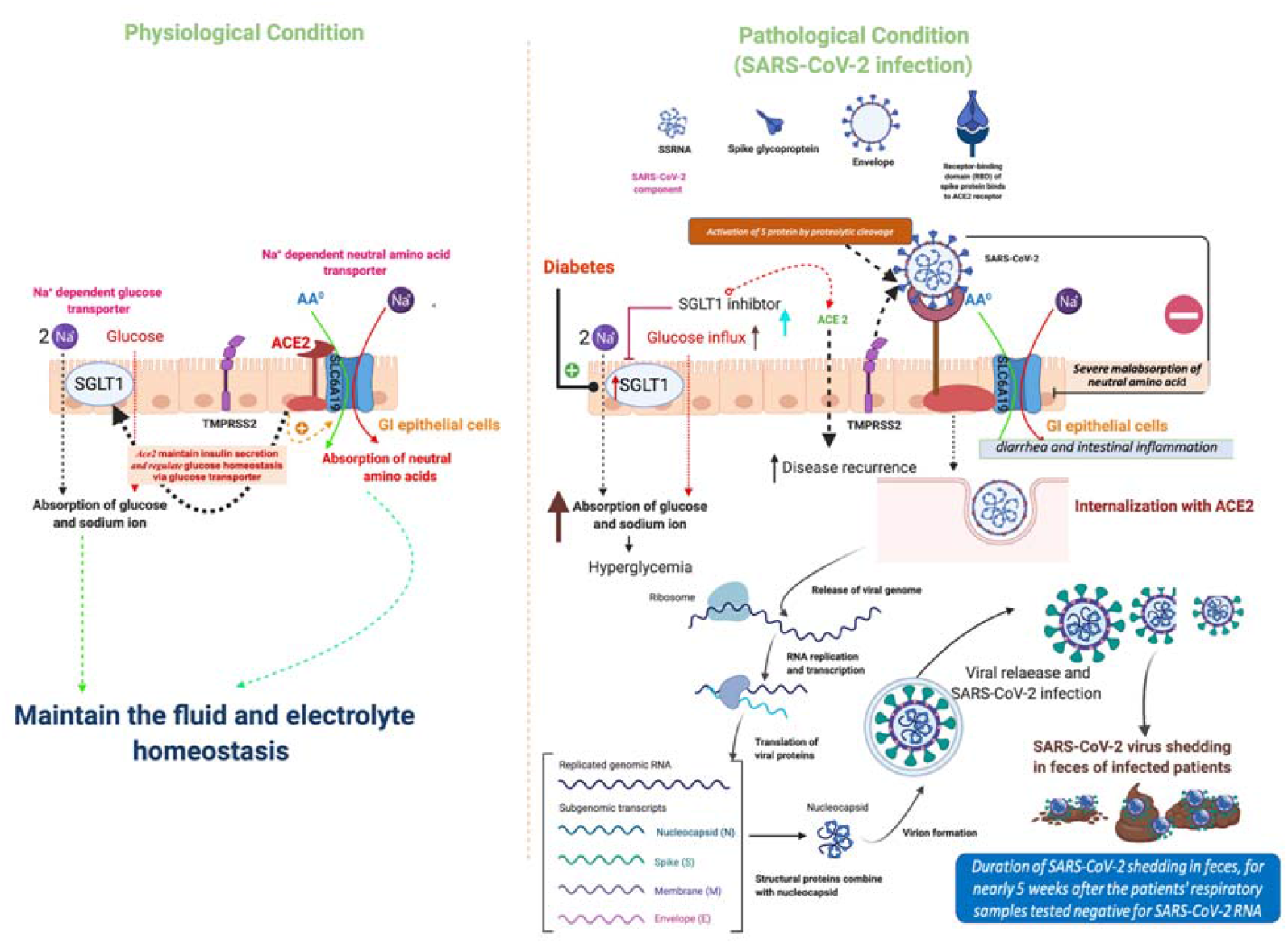

## Introduction

The world is currently reeling in an alarming outbreak of novel coronavirus disease 2019 referred to as COVID-19. COVID-19 is caused by a new coronavirus strain severe acute respiratory syndrome coronavirus 2 (SARS CoV-2)—a positive sense single strand RNA virus. Recent studies which decoded structure of the virus showed binding of its S-spike protein to a human protein-angiotensin converting enzyme 2 (ACE2) (1–3). Following ACE2 binding, cleavage of the viral spike protein (S) by the serine proteases like transmembrane serine protease 2 (TMPRSS2) or Cathepsin L (CTSL) is essential to effectuate host cell membrane fusion and virus infectivity (4). Clinical presentation in COVID-19 patients is highly diverse and majority of them primarily presents with pulmonary symptoms (cough, fever, shortness of breath) (5). In addition, some of the patients present with digestive symptoms like diarrhea, nausea, vomiting and abdominal pain (data ranges from 3.8% to 50.5%) (6). Digestive symptoms have been the only presentations in some of the patients (8,9). Digestive symptoms are not unique to the COVID-19 and usually present in the gastroenteritis caused by many other respiratory syndrome viruses like SARS-CoV-1 and influenza A and B (10,11). However, how SARS-CoV-2 makes entry into the gastrointestinal (GI) tissue leading to gastroenteritis-like features, does not imbibe sufficient and coherent explanation in the light of the existing literature. Some investigators have speculated a fecal-oral route of transmission based on fecal shedding of viral proteins and infectious virus in some COVID-19 patients (12,13).

Knowing the expression pattern of ACE2 and one of the proteases, TMPRSS2 in gastrointestinal tract (GIT) may explicate the pathogenesis of digestive symptoms in COVID-19. Digestive juices and enzymes secreted from the liver, gall bladder (GB) and pancreas play an important role in maintenance of the secretions and absorption of nutrients across intestinal epithelium. Hence their possible dysfunction in COVID-19 patients needs to be examined in order to understand pathogenesis of the digestive symptoms which, in turn, prevent some COVID-19 associated mortality.

Existing literature on the role of ACE2 in regulation of the ion transporters which maintain secretion/absorption across intestinal epithelium provide a clue that digestive symptoms in COVID-19 may have an ACE2 based etiogenesis (11,14–16). Investigating the ACE2 expression pattern of digestive system components may also help to explain exacerbated diabetic complications and mortality in COVID-19 patients. Diabetes has been noted as a co-morbidity (16.2%) in COVID-19 and has contributed to increased mortality (22%) (17) Existing literature implicates ACE2 mediated dysregulation of sodium dependent glucose transporter (SGLT1 or SLC5A1) at intestinal epithelium in the pathogenesis of the diabetes mellitus (18,19).

In this study, we aim at examining the plausibility (based on the tissue specific expression of ACE2) whether any of the digestive system components can be involved in the continued replication of the SARS-CoV-2 after pulmonary symptoms are relieved. Many incidences of disease recurrence have been reported in COVID-19 patients even after being discharged from the hospital. Studies have reported continued shedding of SARS-CoV-2 in the feces of COVID-19 patients up to five weeks after disappearance of the pulmonary symptoms bolstering the indication that a residual persisting of virus inside the digestive system components may be a reason for the disease recurrence (20).

We aimed to validate transcriptomic and proteomic expression of ACE2 and TMPRSS2 in the components of human digestive system (including liver, GB, and pancreas) in tissues derived from the healthy individuals to understand pathophysiological basis of the digestive symptoms in COVID-19 patients.

## Materials and Methods

We analyzed the tissue specific distribution of ACE2 and TMPRSS2 (mRNA and protein) in digestive system components (GIT, liver & GB, and pancreas) using RNA sequencing and immunohistochemistry (IHC) data available in Human Protein Atlas (https://www.proteinatlas.org/humanproteome/sars-cov-2). A digestive system specific functional enrichment map of ACE2 gene was constructed using g:profiler (https://biit.cs.ut.ee/gprofiler/gost) utility and viewed with Cytoscape software, version 3.7.2 (https://cytoscape.org/). Since no direct subject or patient data were used in this study, clearance from the Institutional Ethics Committee was precluded.

### Human Protein Atlas methods

Estimation of mRNA expression and localization of human proteins were performed by the source laboratory using deep sequencing of RNA (RNA-seq) and IHC in normal tissue.

### IHC

As described by the source labs, specimens containing normal tissue were collected and sampled from anonymized paraffin embedded material of surgical specimens, in accordance with approval from the local ethics committee. The specimens were derived from surgical material, normal was defined by morphological parameters and absence of neoplasia. IHC staining was performed using a standard protocol on normal tissue microarray (https://www.proteinatlas.org/download/IHC_protocol.pdf). Antibodies against human ACE2 (HPA000288, CAB026174) and TMPRSS2 (HPA035787) were labeled with DAB (3, 3’-diaminobenzidine) stain. Protein expression score was done based on the staining intensity (negative, weak, moderate or strong) and fraction of stained cells (<25%, 25-75% or >75%). For each protein, the IHC staining profile was matched with mRNA expression data and gene/protein characterization data to yield an ‘annotated protein expression’ profile.

### Transcriptomics

The Human Protein Atlas collects transcriptomic data from the three databases (HPA, GTEx and FANTOM5). HPA RNAseq was performed on human tissue samples from healthy individuals (Accession no: PRJEB4337, Ensembl: ENSG00000130234 (version 92.38). Total RNA was extracted from the tissue samples using the RNeasy Mini Kit (Qiagen, Hilden, Germany) according to the manufacturer’s instructions. The extracted RNA samples were analyzed using either an Experion automated electrophoresis system (Bio-Rad Laboratories, Hercules, CA, USA) with the standard-sensitivity RNA chip or an Agilent 2100 Bioanalyzer system (Agilent Biotechnologies, Palo Alto, USA) with the RNA 6000 Nano Labchip Kit. Only samples of high-quality RNA (RNA Integrity Number 7.5) were used for the mRNA sample preparation for sequencing. mRNA sequencing was performed on Illumina HiSeq2000 and 2500 machines (Illumina, San Diego, CA, USA) using the standard Illumina RNA-seq protocol with a read length of 2×100 bases. Transcript abundance estimation was performed using Kallisto v0.43.1 (https://pachterlab.github.io/kallisto/about). The normalized Tags Per Million (TPM) for each gene from the three databases were calculated and included in the Human Protein Atlas. Each tissue was categorized for the intensity of gene expression using a cutoff value of 1 NX as a limit for detection across all tissues. A tissue was categorized (i) enriched if it had NX level at least four times higher than other tissues, (ii) low specificity if NX ≥ 1 in at least one tissue, (iii) Not detected if NX < 1 in all tissues. Further details of the assays and annotation used by the Human Protein Atlas can be accessed at: https://www.proteinatlas.org/about/assays+annotation#ihk.

### Gene enrichment analysis and visualization

Functional enrichment analysis of the ACE2 gene was performed with g: profiler web server (https://biit.cs.ut.ee/gprofiler/gost) and p-value computed using a Fisher’s exact test with multiple-test correction. Enrichment map visualization was done with the help of Cytoscape software, version 3.7.2 (https://cytoscape.org/).

## Results (Fig. 1-3, S1-3, Table 1, S1-2)

**Figure 1.**
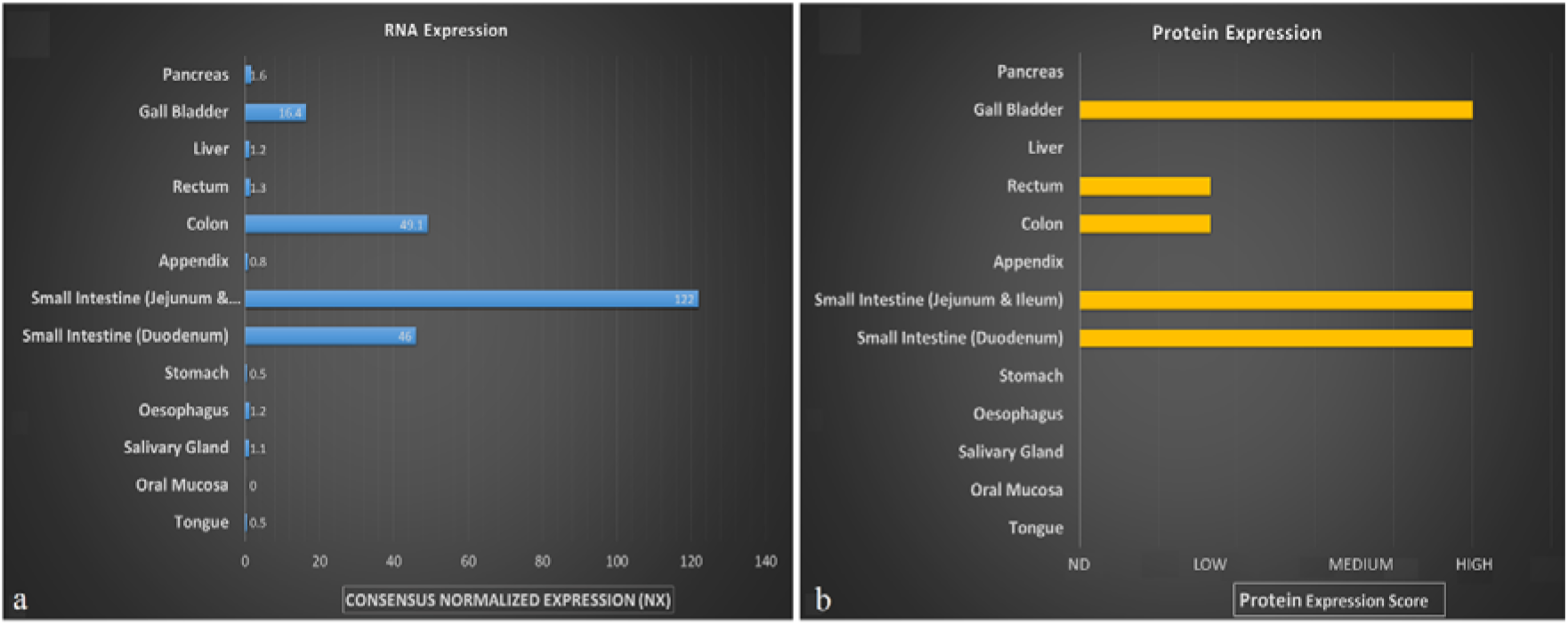
Physiological expression of SARS-CoV-2 binding receptor ACE2 in human digestive system a. mRNA b. Protein. Data Source: The Human Protein Atlas.

**Table 1.** Physiological expression (mRNA and protein) of SARS-CoV-2 binding receptor ACE2 in human digestive system.

| **Tissue** | **Cellular components** | **RNA Expression (NX)** | **Protein Expression** |
| --- | --- | --- | --- |
| Tongue | Squamous epithelial cells | 0.5 | Not detected |
| Oral mucosa | Squamous epithelial cells | 0 | Not detected |
| Salivary gland | Glandular cells | 1.1 | Not detected |
| Esophagus | Squamous epithelial cells | 1.2 | Not detected |
| Stomach | Glandular cells | 0.5 | Not detected |
| Small Intestine<br>(Duodenum) | Glandular cells | 46.0 | High |
| Small Intestine<br>(Jejunum&Ileum) | Glandular cells | 122.0 | High |
| Appendix | Glandular cells | 0.8 | Not detected |
|  | Lymphoid tissue |  | Not detected |
| Colon | Endothelial cells | 49.1 | Not detected |
|  | Glandular cells |  | Low |
|  | Peripheral nerve/ganglion |  | Not detected |
| Rectum | Glandular cells | 1.3 | Low |
| Liver | Bile duct cells | 1.2 | Not detected |
|  | Hepatocytes |  | Not detected |
| Gall bladder | Glandular cells | 16.4 | High |
| Pancreas | Exocrine glandular cells | 1.6 | Not detected |
|  | Islets of Langerhans |  | Not detected |

The transcriptomic and proteomic expression of ACE2 displayed high enrichment in the lower GIT (small intestine, colon, and rectum) (Fig. 1, 2e-h, Table 1). It was highest in the parts of small intestine followed by the colon and the rectum, and nearly absent (negligible/low mRNA expression and undetectable protein expression) in the upper GIT components: mouth cavity (including tongue, oral mucosa, and salivary glands), esophagus, and stomach (Fig. 1, 2a-d). GB showed high glandular expression of ACE2, while any protein expression was undetectable in appendix, liver (hepatocytes and bile duct), and pancreas (exocrine and endocrine glandular tissue) (though minimal mRNA expression was noted) (Fig. 3). Intense ACE2 expression was noted in the glandular cells as well as in the enterocytes in the lining epithelium of the lower GIT (Fig. 2e-h). The cellular expression of ACE2 was visible in the enterocyte cytoplasm and in the apical brush border (Fig. 2e-h, marked with arrow heads). The digestive system specific functional enrichment map for ACE2 gene were related to digestive functions like enzyme activity, amino acids transport, and peptide metabolism at the brush border membrane of enterocytes in the intestinal epithelium (Fig. S1, Table S1). TMPRSS2 was found enhanced in GIT and exocrine glands of pancreas (Fig. S2, Table S2) and found co-localized with ACE2 in enterocytes (Fig. S3).

**Figure 2.**
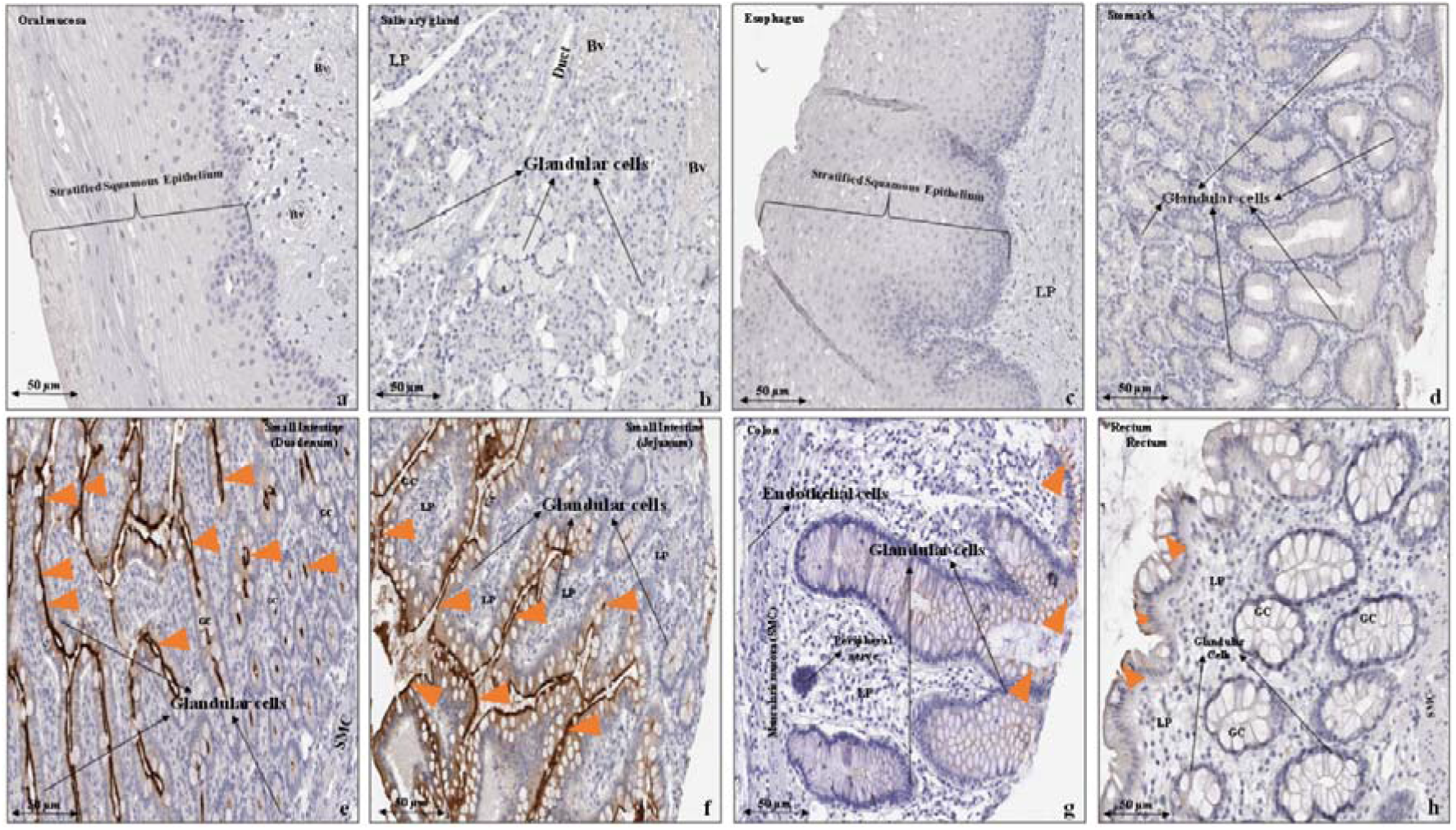
Immunohistochemical expression of ACE2 protein in human gastrointestinal tract a. Oral mucosa b. Salivary gland c. Esophagus d. Stomach e. Duodenum f. Small intestine g. Colon h. Rectum. Orange arrow heads show antibody stained cells. Data Source: The Human Protein Atlas. **Abbreviations:** GC- goblet cells, Bv - Blood vessels, LP - Lamina propria, SMC - Smooth muscle cells.

**Figure 3.**
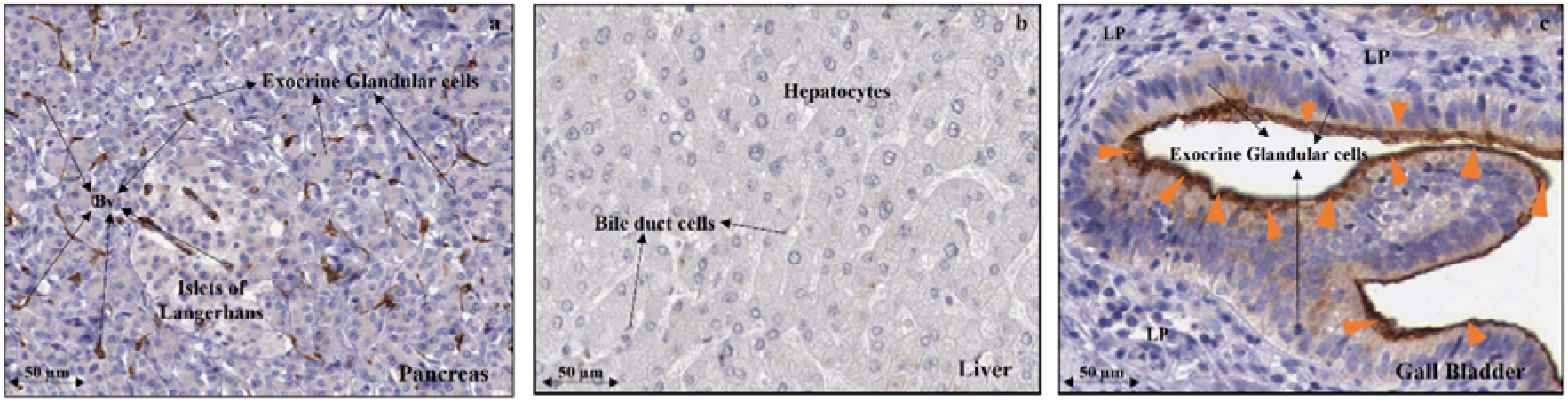
Immunohistochemical expression of ACE2 protein in Human tissue a. Pancreas b. Liver c. Gall bladder. Orange arrow heads show antibody stained cells. (In pancreatic tissue blood vessels (Bv) but not in the exocrine or endocrine glandular cells can be seen expressing ACE2.) Data Source: The Human Protein Atlas. **Abbreviations:** Bv - Blood vessels, LP - Lamina propria, SMC - Smooth muscle cells.

## Discussion

We found enriched transcriptomic and proteomic expression of SARS-CoV-2 binding receptor ACE2 in lower GIT (small intestine, colon, and rectum) and GB (Fig. 1-3. Table 1). The digestive system specific functional enrichment map of the ACE2 gene suggests its role in regulating secretory/absorptive functions at the brush border membrane of the enterocytes in the intestinal lining epithelium (Fig.S1, Table S1). The co-localized expression of SARS-CoV-2 cell entry associated protease TMPRSS2 in the enterocytes make these cells potential sites for viral infection (Fig. S2-3, Table S2).

ACE2 is a homologue of angiotensin-I converting enzyme (ACE), the key enzyme of the renin-angiotensin system (RAS). It is an integral membrane protein and localizes predominantly at the apical surface of polarized epithelial cells where it is proteolytically cleaved within its ectodomain to release a soluble form (21,22). Currently, SARS-CoV-2 mediated binding of ACE2 and the following downstream events leading to tissue damage are little known. Presumptive understanding of SARS-CoV-2 driven pathology is being borrowed from SARS-CoV-1 which was the etiological basis of SARS pandemic in 2003. Uniquely, it acted on the same receptor as SARS-CoV-2 and led to many clinical manifestations similar to COVID-19 (23). Studies utilizing cell lines to decipher SARS pathology in lung tissue showed that the spike protein of SARS-CoV-1 (SARS-S) induced TNFα production which facilitated virus entry (24). TNFα also led to inflammation of the cell membrane and consequently tissue damage (22–24). SARS-CoV-1 was also showed to cause downregulation of ACE2 expression at the cell membrane level (22,25). Existing literature regarding expression of ACE2 in human tissues are rare. Hamming et al, studied ACE2 protein expression in human tissues in reference to SARS-CoV-1 (26). Our findings for ACE2 protein expression in digestive system components are in line with the findings of their study (26). Recently, enriched expressions of ACE2 (and TMPRSS2) in enterocytes and mucus producing cells were shown using single cell m-RNA expression studies (27,28). Enriched expression of SARS-CoV-2 binding receptor ACE2 in the mucosal glands and enterocytes (including brush border cells) in the lining epithelium (Fig. 2e-h, Table 1) of the lower GIT indicates that GI cells are potential sites for virus replication. Evidence of the viral shedding in the feces shown in some studies indicates possible replication of the virus inside the GI cells which, in turn may explain GI manifestations of COVID-19 in addition to disease recurrence (29,30). Recent *in situ* studies using recombinant strain of SARS-CoV-2 showed that the virus can potentially infect and replicate in human intestinal tissue (31,32). Further, GIT to pulmonary spread of SARS-CoV-2 infection has been indicated by a study by Sun *et al* who showed in a transgenic mouse expressing human ACE2 that a direct intragastric inoculation of SARS-CoV-2 can cause productive infection and lead to pulmonary pathological changes (33).

How the virus reaches the GI is arguable? Some authors speculated a fecal-oral route of entry (8). Shedding of infectious SARS-CoV-2 in feces was also detected in occasional COVID-19 patients (12,13). We examined possibility of this route of entry based on the expression pattern of ACE2 along the length of the GIT (Fig. 1, 2, Table 1). Negligible or very low mRNA expression and undetectable proteomic expression of ACE2 in the mouth cavity (including tongue, oral mucosa, and salivary glands), esophagus, and stomach (Fig. 1, 2a-d, Table 1) indicate these parts of GIT can be resistant for the virus entry. But this observation does not negate a possible site of virus entry through the ACE2 receptors present in the lower GIT in case of fecal-oral transmission. It is then intriguing that how SARS-CoV-2 survives extremes of pH within the digestive system milieu (gastric-1.5 to 3.5, pancreatic-7.5, bile acid-7-8) while passing along the length of GIT. Recently, Chin *et al.,* 2020 showed *in vitro* that SARS-CoV-2 can survive at wide range of pH values at room temperature (pH3-10) (34). This can be further explained by an earlier study by Hirose et al, who, in an experimental, model demonstrated that RNA viruses like influenza A and B (when swallowed) can survive extremes of pH and maintain infectivity with help of the mucus cover lining GIT allowing their safe passage and even excretion in feces (35). Mucus cells are abundant all along the length of the GIT which can contribute to the carriage and survival of SARS-CoV-2 thereby contributing to the so hypothesized fecal-oral transmission. This also hints that shedding of the virus in feces always may not be indicative of its replication in GI cells; all those patients who shed virus in stools don’t necessarily present with digestive symptoms (29).

Healthy intestinal mucosa may not be well conducive for the entry of the virus due to the presence of unique multi-layer barrier system, though a prior inflammatory condition which disrupts mucosal barrier may render the lower GI entry of the SARS-CoV-2 using ACE2 receptor and its replication inside tissue plausible (36). Inflammatory conditions in GIT enhance the expression of ACE2 in the luminal epithelium which can provide additional support for the entry of the virus (37). Once inside the GI cells, the virus can replicate there and may orchestrate viral toxin mediated cell injury ensuing further inflammation, thereby, giving rise to gastroenteritis like symptoms (diarrhea, nausea, and vomiting, abdominal pain) (22,24,38). Other than the fecal-oral route, an alternative route of viral entry to the GI cells may be through the tissue microvasculature. Though this may not be highly probable but this premise does warrant consideration. In that case, fecal viral shedding can happen after sloughing of the inflamed/necrosed intestinal mucosa. Currently, data is limited which support presence of SARS-CoV-2 in the blood, however such evidence is available for other coronaviruses infections like SARS and MERS (29,39–41).

ACE2 is known to regulate sodium-dependent amino acid and glucose transporters in the enterocytes brush border which physiologically engage in the absorption of nutrients from the digested food, and maintain osmotic and electrolyte balance across the GI lining epithelium (11,14). In a recent study Yan et al., 2020 showed that SARS-CoV-2 can bind to the complex of ACE2 with B0AT1(Slc6a19)—a major sodium dependent neutral amino acid transporter present in the epithelial lining of human intestine (and also in kidneys) (1,42). The dysregulation of the intestinal ion transporters has been implicated in the pathophysiology of infectious diarrhea and malabsorption disorders (15,16). Literature also suggests that a dysregulation of these transporters can ensue interleukin/cytokine mediated intestinal inflammation and can give rise to digestive symptoms (14). An enhanced GI expression of ACE2 is known in inflammatory bowel diseases (IBDs) which present with similar symptoms as in COVID-19 patients (14,43).

Based on the findings of this study and supportive evidence from the literature, we propose that a virus binding-ACE2 mediated dysregulation of the sodium dependent nutrient transporters may be a plausible basis for the digestive symptoms in COVID-19. Prior intestinal inflammatory conditions like IBD may raise the susceptibility of SARS-CoV-2 infection through fecal-oral transmission. ACE2 mediated dysregulation of SGLT1 and/or SLC5A1 at intestinal epithelium also links it to the pathogenesis of diabetes mellitus (18,19). The SGLT1 transporters are physiologically involved in active absorption of glucose across the intestinal epithelium and its virus binding receptor ACE2 mediated dysregulation may exacerbate the existing impaired glycemic control in COVID-19 patients with diabetes mellitus (19). (Sufficient data on glycemic control in COVID-19 patients is lacking for now, impaired glycemic control was stated as an independent risk factor predicting morbidity and mortality in SARS patients with diabetes mellitus (44).) ACE2 mediated downregulation of SGLT1 in intestinal epithelium prevents hyperglycemia in rat models of the diabetes mellitus (45,46). Though direct evidence is lacking in terms of the effect of SARS-CoV-2 binding on ACE2 on its signaling cascades, however, substantiation from SARS-CoV-1 studies (for SARS) suggests that it can downregulate ACE2 expression (25). Such an eventuality can lead to upregulation of SGLT1 thereby precipitating hyperglycemia (45,46). (SGLT1 inhibitors are being used in treatment of diabetes mellitus, their use in COVID-19 patients may need a rethinking for the dose adjustments (47).)

Our data showed undetectable expression of ACE2 and TMPRSS2 proteins in insulin producing Islets of Langerhans of the pancreas raising an insulin independent possibility of dysregulated intestinal SGLT1 transporters. This bolsters the rationale behind diabetes related increased morbidity/mortality in COVID-19 patients. Apart from intestine SGLT1 is known to be widely expressed in other human tissues like proximal tubule of kidney, heart, and liver (proteinatlas.org/ENSG00000100170-SLC5A1/tissue) where it regulates the glucose absorption. An ACE2-mediated dysregulation of SGLT1 in COVID-19 patients warrants further investigation.

High expression of ACE2 in glandular cells of the GB indicates that this also can be a potential site for the virus replication. (Contrastingly, we found low m-RNA and undetectable proteomic expression of TMPRSS2 in glandular cells of GB, however, robust expression of another serine protease CTSL is noted in these cells in the records of Human Protein Atlas (48), which may be able to substitute for TMPRSS2 (1)) GB has a luminal connection to the duodenum through cystic and common bile duct (CBD). Though this connection is guarded by a sphincter (of Oddi) present in duodenal mucosa, it doesn’t create an anatomical barrier and, therefore, a viral invasion along the mucosal epithelium remains a possibility.

GB is the physiological storage site for the bile secreted from the hepatocytes, and pathology of this organ can also contribute to the digestive symptoms present in COVID-19 patients. GB has been a known reservoir for *Salmonella typhi,* a bacterium causing enteric fever, and one of the cited reasons for disease recurrence (49). The thick mucin secreted from its glandular cells can provide a protective environment for survival of SARS-CoV-2 (as we discussed above for GI lining epithelium) (35). Hence, GB homing may act as a mechanism for the replication of the virus even without ensuing a local tissue injury.

Continued replication of the SARS-CoV-2 in the intestinal tissue, and possibly in GB, may be a potential reason for the recurrence of SARS-CoV-2 in the light of the diagnostic tests as has been noted in some COVID-19 patients after being discharged from the hospital (40,50). A post-mortem study of these organs in COVID-19 patients may provide some confirmation in this regard.

Based on the observed pattern of tissue specific expression of ACE2 (which binds to SARS-CoV-2) in the components of the digestive system in normal individuals, we propose that an ACE2 based mechanism may be involved in the pathogenesis of digestive symptoms, increased diabetes-associated mortality risk, and disease recurrence in COVID-19.

### Limitations

All the aspects of the plausible SARS-CoV-2 binding receptor ACE2 mediated pathology in the digestive system which we have discussed above are based on the distribution of the virus cell entry related factors in the normal tissue. Hence, this study presents indirect evidence which needs to be validated in actual patients before reaching any conclusion.

### Future directions

Further studies are advisable to understand the molecular mechanisms involved in the SARS-CoV-2 binding receptor ACE2 mediated dysregulation of the intestinal nutrient transporters and finding out COVID-19 specific drug targets. Inter-individual variations in frequency of the digestive symptoms, diabetes associated mortality, and recurrences may depend upon the genotype specific variations in ACE2 expression and other patient specific characteristics (like age, sex, and comorbidity). A study of these variables in the disease pathogenesis may help in deciding personalized therapeutic management for the COVID-19 cases.

## Author Contributions

AK conceived the idea. AK wrote the first draft. MAF, VP, KR, MK, CK, KK, PK, PP, HN, RKN, SNP, RQ, and SK revised the draft. RKN, KR, PP, PK, and VP contributed to data analysis, and prepared tables and figures.

## Data Availability

Data used for this study can be accessed at the following link: https://www.proteinatlas.org/about/download

## Acknowledgments

The authors acknowledge The Human Protein Atlas (https://www.proteinatlas.org/) for ready availability of data in the public domain. This manuscript has been released as a pre-print at BioRxiv [bioRxiv 2020.04.14.040204, Kumar A, et al. (51)].

---

**Figure S1.**
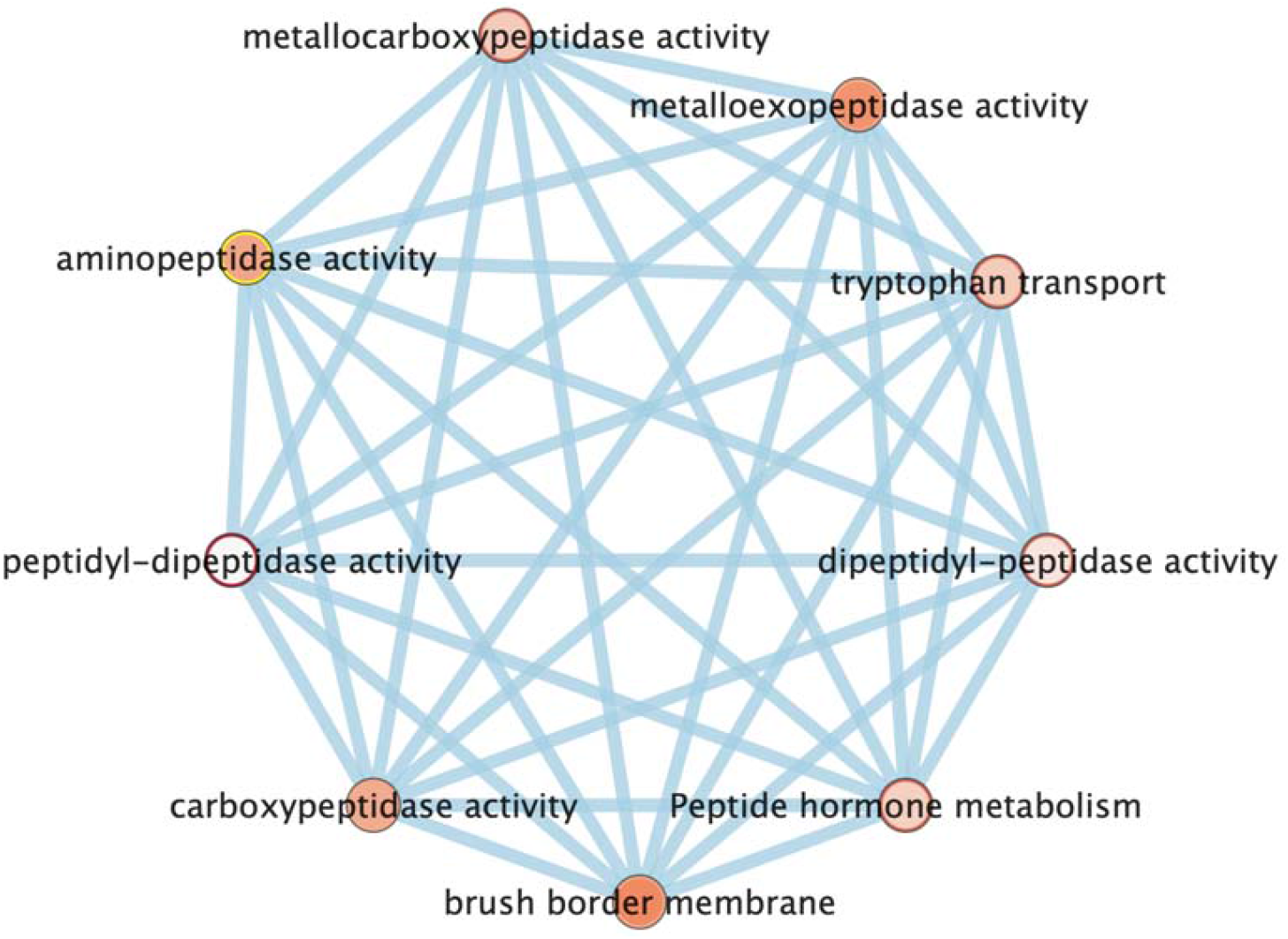
ACE2 gene enrichment map for Digestive system functions.

**Figure S2.**
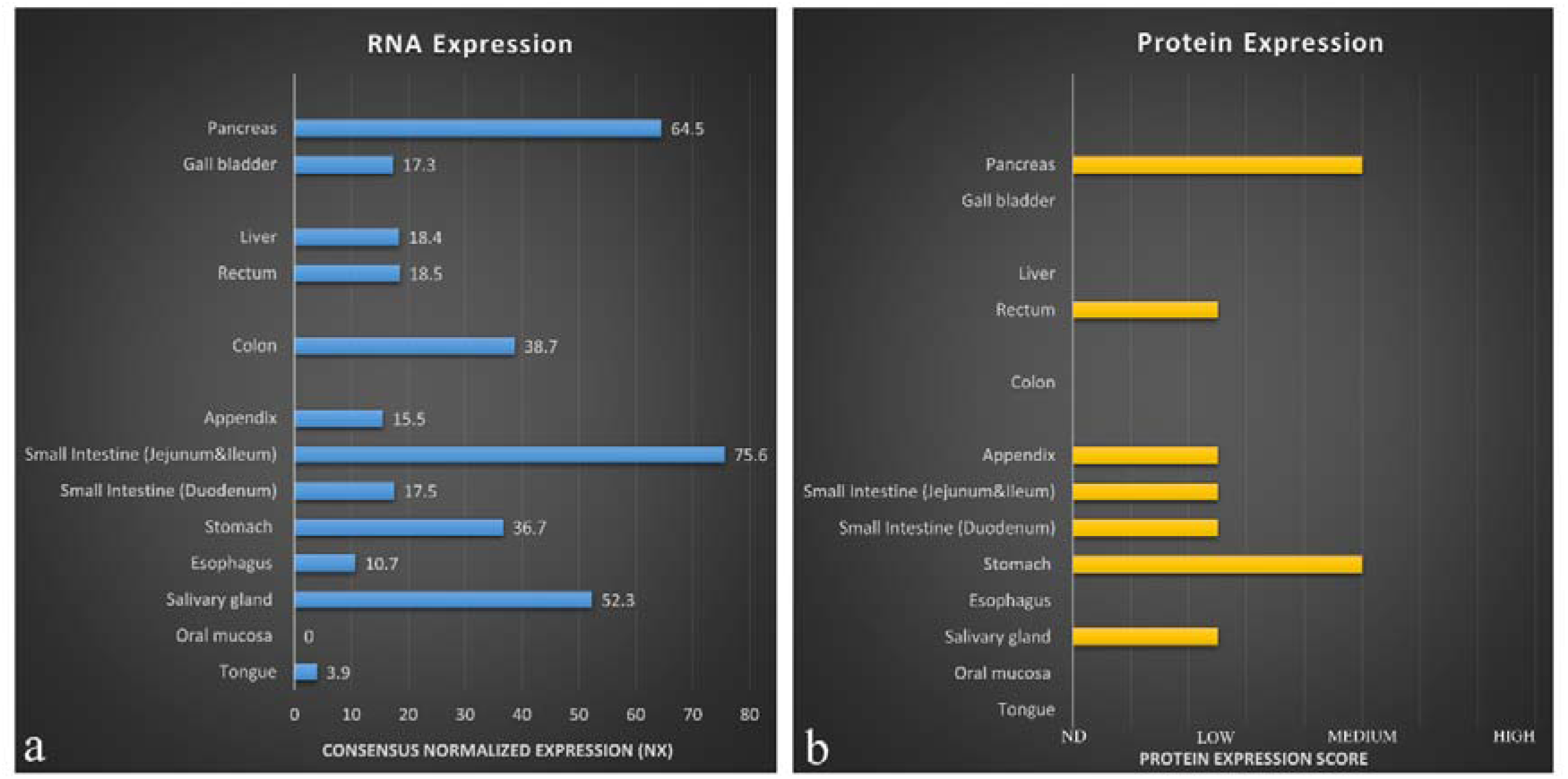
Physiological expression of SARS-CoV-2 cell entry associated protease TMPRSS2 in human digestive system a. mRNA b. Protein. Data Source: The Human Protein Atlas.

**Figure S3.**
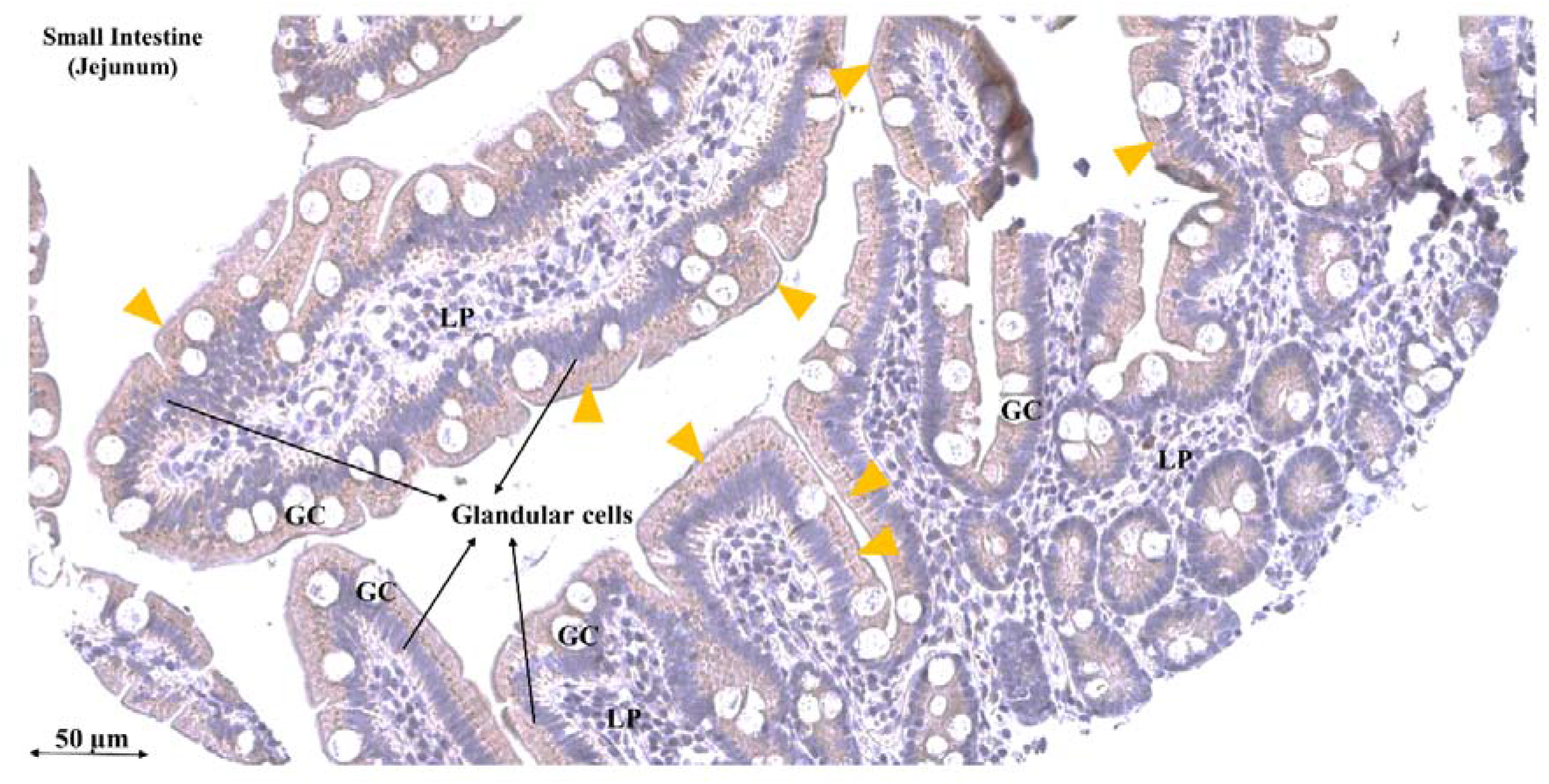
Immunohistochemical expression of TMPRSS2 protein in Small Intestine of human gastrointestinal tract Orange arrow heads show antibody stained cells. Data Source: The Human Protein Atlas. **Abbreviations:** GC- goblet cells, LP - Lamina propria.

**Table S1.** ACE2 gene enrichment for Digestive system functions.

| **GO. ID** | **Description** | **P Value** | **FDR** | **Phenotype** | **Gene** |
| --- | --- | --- | --- | --- | --- |
| GO:0008241 | Peptidyl-dipeptidase activity | 0.00397219 | 0.00397219 | 1 | ACE2 |
| GO:0008239 | Dipeptidyl-peptidase activity | 0.01853691 | 0.01853691 | 1 | ACE2 |
| GO:0004181 | Metallocarboxypeptidase activity | 0.03707382 | 0.03707382 | 1 | ACE2 |
| GO:0004180 | Carboxypeptidase activity | 0.06090698 | 0.06090698 | 1 | ACE2 |
| GO:0004177 | Aminopeptidase activity | 0.06487918 | 0.06487918 | 1 | ACE2 |
| GO:0008235 | Metalloexopeptidase activity | 0.07944389 | 0.07944389 | 1 | ACE2 |
| GO:0140272 | Exogenous protein binding | 0.10062893 | 0.10062893 | 1 | ACE2 |
| GO:0008238 | Exopeptidase activity | 0.1509434 | 0.1509434 | 1 | ACE2 |
| GO:0008237 | Metallopeptidase activity | 0.23965574 | 0.23965574 | 1 | ACE2 |
| GO:0004175 | Endopeptidase activity | 0.58126448 | 0.58126448 | 1 | ACE2 |
| GO:0070011 | Peptidase activity, acting on L-amino acid peptides | 0.8142999 | 0.8142999 | 1 | ACE2 |
| GO:0008233 | Peptidase activity | 0.84872559 | 0.84872559 | 1 | ACE2 |
| GO:0005515 | Protein binding | 1 | 1 | 1 | ACE2 |
| GO:0140096 | Catalytic activity, acting on a protein | 1 | 1 | 1 | ACE2 |
| GO:0015827 | Tryptophan transport | 0.03708254 | 0.03708254 | 1 | ACE2 |
| GO:0051957 | Positive regulation of amino acid transport | 0.17614208 | 0.17614208 | 1 | ACE2 |
| GO:0032800 | Receptor biosynthetic process | 0.24103652 | 0.24103652 | 1 | ACE2 |
| GO:0003081 | Regulation of systemic arterial blood pressure by renin-angiotensin | 0.2595778 | 0.2595778 | 1 | ACE2 |
| GO:0051955 | Regulation of amino acid transport | 0.30593097 | 0.30593097 | 1 | ACE2 |
| GO:0016486 | Peptide hormone processing | 0.32447224 | 0.32447224 | 1 | ACE2 |
| GO:1901890 | Positive regulation of cell junction assembly | 0.33374288 | 0.33374288 | 1 | ACE2 |
| GO:0032892 | Positive regulation of organic acid transport | 0.33374288 | 0.33374288 | 1 | ACE2 |
| GO:0051954 | Positive regulation of amine transport | 0.34301352 | 0.34301352 | 1 | ACE2 |
| GO:1903793 | Positive regulation of anion transport | 0.49134368 | 0.49134368 | 1 | ACE2 |
| GO:0051952 | Regulation of amine transport | 0.87143974 | 0.87143974 | 1 | ACE2 |
| GO:0044070 | Regulation of anion transport | 0.91779292 | 0.91779292 | 1 | ACE2 |
| GO:0019538 | Protein metabolic process | 1 | 1 | 1 | ACE2 |
| GO:0019222 | Regulation of metabolic process | 1 | 1 | 1 | ACE2 |
| GO:0015849 | Organic acid transport | 1 | 1 | 1 | ACE2 |
| GO:0015711 | Organic anion transport | 1 | 1 | 1 | ACE2 |
| GO:0010817 | Regulation of hormone levels | 1 | 1 | 1 | ACE2 |
| GO:0009893 | Positive regulation of metabolic process | 1 | 1 | 1 | ACE2 |
| GO:0016485 | Protein processing | 1 | 1 | 1 | ACE2 |
| GO:0032879 | Regulation of localization | 1 | 1 | 1 | ACE2 |
| GO:0046942 | Carboxylic acid transport | 1 | 1 | 1 | ACE2 |
| GO:0043270 | Positive regulation of ion transport | 1 | 1 | 1 | ACE2 |
| GO:0043269 | Regulation of ion transport | 1 | 1 | 1 | ACE2 |
| GO:0043170 | Macromolecule metabolic process | 1 | 1 | 1 | ACE2 |
| GO:0043112 | Receptor metabolic process | 1 | 1 | 1 | ACE2 |
| GO:0042445 | Hormone metabolic process | 1 | 1 | 1 | ACE2 |
| GO:0008152 | Metabolic process | 1 | 1 | 1 | ACE2 |
| GO:0001816 | Cytokine production | 1 | 1 | 1 | ACE2 |
| GO:0001817 | Regulation of cytokine production | 1 | 1 | 1 | ACE2 |
| GO:0006820 | Anion transport | 1 | 1 | 1 | ACE2 |
| GO:0006812 | Cation transport | 1 | 1 | 1 | ACE2 |
| GO:0006811 | Ion transport | 1 | 1 | 1 | ACE2 |
| GO:0006807 | Nitrogen compound metabolic process | 1 | 1 | 1 | ACE2 |
| GO:0006518 | Peptide metabolic process | 1 | 1 | 1 | ACE2 |
| GO:0006508 | Proteolysis | 1 | 1 | 1 | ACE2 |
| GO:0071705 | Nitrogen compound transport | 1 | 1 | 1 | ACE2 |
| GO:0071704 | Organic substance metabolic process | 1 | 1 | 1 | ACE2 |
| GO:0071702 | Organic substance transport | 1 | 1 | 1 | ACE2 |
| GO:0051604 | Protein maturation | 1 | 1 | 1 | ACE2 |
| GO:0051050 | Positive regulation of transport | 1 | 1 | 1 | ACE2 |
| GO:0051049 | Regulation of transport | 1 | 1 | 1 | ACE2 |
| GO:0031526 | Brush border membrane | 0.08612643 | 0.08612643 | 1 | ACE2 |
| GO:0005903 | Brush border | 0.16610098 | 0.16610098 | 1 | ACE2 |
| REAC:R-HSA-2980736 | Peptide hormone metabolism | 0.03360393 | 0.03360393 | 1 | ACE2 |
| REAC:R-HSA-392499 | Metabolism of proteins | 0.760808 | 0.760808 | 1 | ACE2 |

**Table S2.** Physiological expression (mRNA and protein) of SARS-CoV-2 cell entry associated protease TMPRSS2 in human digestive system.

| **Tissue** | **Cellular components** | **RNA Expression (NX)** | **Protein Expression** |
| --- | --- | --- | --- |
| Tongue | Squamous epithelial cells | 3.9 | Not detected |
| Oral mucosa | Squamous epithelial cells | 0 | Not detected |
| Salivary gland | Glandular cells | 52.3 | Low |
| Esophagus | Squamous epithelial cells | 10.7 | Not detected |
| Stomach | Glandular cells | 36.7 | Medium |
| Small Intestine (Duodenum) | Glandular cells | 17.5 | Low |
| Small Intestine (Jejunum & Ileum) | Glandular cells | 75.6 | Low |
| Appendix | Glandular cells | 15.5 | Low |
|  | Lymphoid Tissue |  | Not detected |
| Colon | Endothelia cells | 38.7 | Not detected |
|  | Glandular cells |  | Not detected |
| Rectum | Glandular cells | 18.5 | Low |
| Liver | Bile duct cells | 18.4 | Not detected |
|  | Hepatocytes |  | Not detected |
| Gall bladder | Glandular cells | 17.3 | Not detected |
| Pancreas | Exocrine glandular cells | 64.5 | Medium |
|  | Islets of Langerhans |  | Not detected |

